# Functional implication of the homotrimeric multidomain vacuolar sorting receptor 1 (VSR1) from *Arabidopsis thaliana*

**DOI:** 10.1101/2023.12.15.571892

**Authors:** HaJeung Park, BuHyun Youn, Daniel J. Park, Sathyanarayanan V. Puthanveettil, ChulHee Kang

## Abstract

The vacuolar sorting receptors (VSRs) are specific to plants and are responsible for sorting and transporting particular proteins from the *trans*-Golgi network to the vacuole. This process is critically important for various cellular functions, including storing nutrients during seed development. Despite many years of intense studies on VSRs, a complete relation between function and structure has not yet been revealed. For the first time, the crystal structure of the full-length luminal part of glycosylated VSR1 from *Arabidopsis thaliana* (AtVSR1) has been determined. The structure provides insights into the tertiary and quaternary structures of VSR1, which are composed of an N-terminal protease-associated (PA) domain, a unique central region, and one epidermal growth factor (EGF) domain followed by two disordered EGF domains. The structure of VSR1 exhibits unique characteristics, the significance of which is yet to be fully understood.

## Introduction

The proper functioning of eukaryotic cells requires the sorting and targeting of proteins synthesized in the endoplasmic reticulum (ER). From the ER, correctly folded/assembled proteins are transferred to the Golgi, which initiates the first step in protein sorting and trafficking. In plant cells, vacuolar sorting receptors (VSRs) are responsible for the sorting of proteins from the *trans*-Golgi network (TGN) to prevacuolar compartments (PVCs) and finally to their respective vacuoles (Kang and Hwang, 2014). VSRs are type-I transmembrane proteins with ∼600 amino acids (80 kDa without the N-terminal signaling peptide) (Paris et al., 1997). VSRs recognize specific signal sequences of the cargo proteins called Vacuole Sorting Determinants (VSDs) through its N-terminal protease-associated (PA) domain (Cao et al., 2000; Luo et al., 2014; Tsao et al., 2022). This recognition by VSRs is known to be pH-dependent; the cargo protein binds to the VSR via the VSD in the TGN in a neutral pH, and the cargo protein is released upon exposure to an acidic pH (Reguera et al., 2015). In addition, PV72, a homolog of VSR in potatoes has a calcium-dependent cargo binding within the pH range of 5.5 to 7 (Watanabe et al., 2002).

The VSDs of various vacuolar proteins are mainly classified into two groups, sequence-specific vacuolar sorting determinants (ssVSDs) (Kirsch et al., 1994, 1996) and C-terminal vacuolar sorting determinants (ctVSDs) (Shimada et al., 2003). The proaleurain peptide (SSSFADSNPIRPVTDRAASTYC) present in a plant cysteine protease is the prime example of a ssVSD with the presence of a central ‘NPIR’ motif (Kirsch et al., 1994, 1996). Any alteration from this specific sequence motif can negatively affect AtVSR1-binding. For instance, a mutated peptide with glycine in place of isoleucine in the NPIR motif renders the peptide unable to compete with the proaleurain peptide for binding to VSR1 (Kirsch et al., 1994, 1996)). On the other hand, the characteristics of ctVSDs appear to be contextual without any distinctive known sequence features, though it has been proposed that the VSR1-binding motif is approximately five residues long, with the last three being hydrophobic, and it is located at the C-terminus of a cargo protein (Shimada et al., 1997; Tsao et al., 2022). A recent study by Park *et al*. proposed that soluble proteins carrying a ctVSD are transported by RMRs, not by AtVSR1 (Park, Oufattole and Rogers, 2007).

The structural details of PA domain of AtVSR1 in complexes with both ssVSD and ctVSD were reported ((Shimada et al., 1997; Tsao et al., 2022; Luo et al., 2014). The structures show that the peptides interact with the PA domain of AtVSR1 in sequence non-specific manners, as the peptides bind the PA domain through their backbone only without any side chain involvement. Furthermore, the PA-ssVSD complex showed that the PA interacted with the preceding residues of the NPIR motif, while the motif itself was invisible, probably because of structural disorder (Luo et al., 2014). Despite these structural studies, the involvement of the PA domain in recognizing VSD motifs remains unclear.

Herein, we report a crystal structure of full-length VSR1 luminal domain from *Arabidopsis thaliana* (AtVSR1). The structure shows a distinct organization of constituting domains with unique interactions among them. Although the structure does not contain a bound peptide, it still offers valuable insights into the ligand-binding mechanism of VSRs. Additionally, the crystal contacts and domain structures offer valuable insights into the other characteristics and potential functions of this unique class of proteins.

## Results

### The overall structure

The luminal part of AtVSR1 from residue 21 to 582 was expressed from *Drosophila* S2 insect cells. The purified protein was crystallized in a space group P2_1_3, which diffracted up to 2.6 Å (Rogers et al., 2004). The crystal lattice packing indicated that one molecule in the asymmetric unit was closely associated with neighboring molecules through crystallographic three-fold symmetry. The resulting crystal structure was composed of an N-terminal protease-associated (PA) domain, a unique central region, and three C-terminal epidermal growth factor (EGF)-like repeats (Cao et al., 2000) (**Fig. 1A**). The resolved structure had disordered regions at the C-terminus, where density could not be uniquely traced for proper model building after the first EGF-like domain. At the beginning of the refinement, a clear electron density attached to the sidechain of Asn289 in the central domain was observed and assigned as two N-acetylglucosamine (GlcNAc).

**Figure 1.**
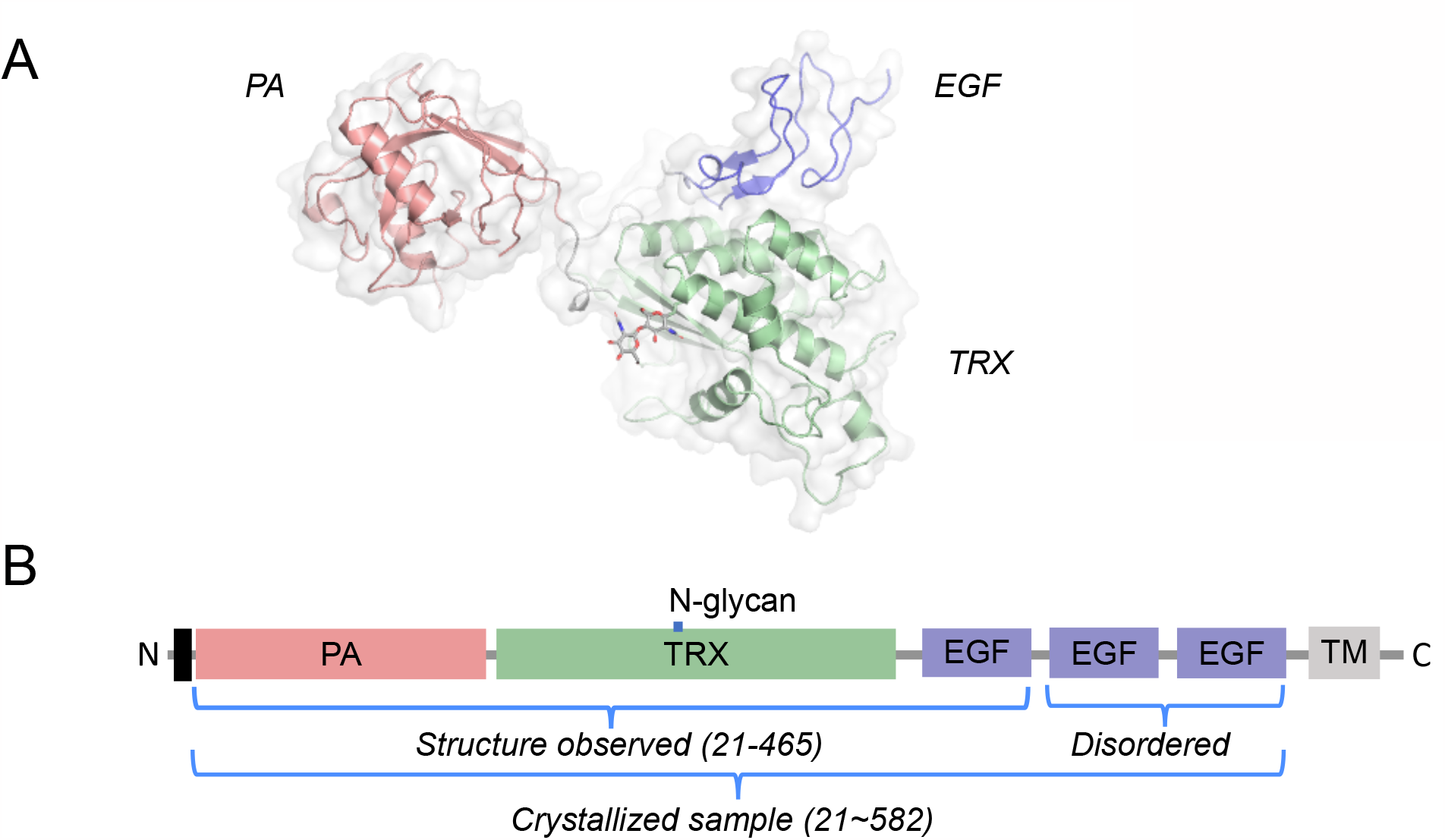
The overall structure and the schematic of AtVSR1. **A**, The overall structure of AtVSR is shown as a cartoon model. The color of each domain matches that of the schematic. Each domain is also labeled. **B**, The schematic of AtVSR1 shows the domain organizations, the information of the crystallization construct, and the structure resolved portion. N-glycan indicates the location where glycosylation is observed. The black rectangle represents the signaling sequence.

The structural model starts at Phe21 and ends at Ala465 and the overall structure is separated into three distinct domains: the PA domain (Phe21-Trp178), the thioredoxin (TRX) domain (Val186-Phe396), and the EGF-like domain (Glu409-Ala462) (**Fig. 1B**). The central domain is named as TRX domain, as a DALI search picked γ-interferon-inducible lysosomal thiol reductase (PDB ID 6NWX) as the closest 3D structure with a high Z-score of 14.6, followed by bacterial disulfide isomerase A (PDB ID 3BCI) with a Z-score of 14.0 (Holm et al., 2023). The rest of the DALI results are all TRX-fold proteins, most of which are involved in protein disulfide bond reduction (**Table 1**).

**Table 1.** Structural similarity search by the Dali server.

| No | PDB ID-Chain | Z | rmsd (Å) | Alignment length | %id | Description | Organism |
| --- | --- | --- | --- | --- | --- | --- | --- |
| 1 | 6nwx-A | 14.6 | 3.0 | 162 | 10 | Gamma-interferon-inducible lysosomal thiol reductase | Mus musculus |
| 2 | 3bci-A | 14.0 | 2.7 | 153 | 14 | DsbA | Staphylococcus aureus |
| 3 | 3bck-A | 13.9 | 2.7 | 152 | 14 | DsbA T153V | Staphylococcus aureus |
| 4 | 3bd2-A | 13.8 | 2.7 | 152 | 14 | DsbA E96Q | Staphylococcus aureus |
| 5 | 2in3-A | 12.6 | 2.9 | 155 | 10 | PDI | Nitrosomonas europaea |
| 6 | 6ghb-D | 12.3 | 2.9 | 159 | 9 | YjbH | Geobacillus kaustophilus |
| 7 | 3eu3-A | 12.2 | 3.3 | 156 | 14 | BdbD | Bacillus subtilis |
| 8 | 3gha-A | 12.1 | 3.2 | 156 | 14 | BdbD | Bacillus subtilis |
| 9 | 3eu4-A | 12.1 | 3.3 | 156 | 14 | BdbD | Bacillus subtilis |
| 10 | 3gh9-A | 12.1 | 3.3 | 156 | 14 | BdbD | Bacillus subtilis |
| 11 | 5vyo-B | 11.6 | 3.5 | 153 | 18 | DsbA | Burkholderia pseudomallei |
| 12 | 4jr4-A | 11.6 | 3.0 | 157 | 13 | DsbA | Mycobacterium tuberculosis |
| 13 | 4jr6-A | 11.6 | 3.0 | 157 | 12 | DsbA | Mycobacterium tuberculosis |
| 14 | 3kzq-B | 11.6 | 3.5 | 159 | 11 | Putative protein | Vibrio parahaemolyticus |
| 15 | 5hfi-A | 11.5 | 2.9 | 154 | 9 | Disulfide reductase DsbM | Pseudomonas aeruginosa |
| 16 | 5vyo-C | 11.5 | 3.5 | 153 | 18 | DsbA | Burkholderia pseudomallei |
| 17 | 4k6x-B | 11.5 | 3.0 | 156 | 13 | DsbA-like oxidases | Mycobacterium tuberculosis |

The PA and the TRX domains are connected by a 7-residue linker with no significant interaction between them. In contrast, the TRX and the EGF-like domains are connected through a 12-residue linker and display tight interdomain interactions between them through an extensive hydrogen bond (H-bond) network (**Suppl. Table 1**). At the core of the interdomain interface, the side chain of Asp303 in TRX forms H-bonds with the side chains of Arg440 and Tyr457 in EGF-like domains (**Fig. 2A and B**). At the periphery, an additional salt bridge between Lys 221 and Glu414 is observed, and the main chain carbonyl of Arg440 also forms an H-bond with the amine side chain of Lys213 (**Fig. 2C**) Notably, no significant hydrophobic interaction was found in this interdomain interaction. The Cys405 in the linker establishes a disulfide bond with Cys393 of the TRX domain potentially stabilizing the linker.

**Figure 2.**
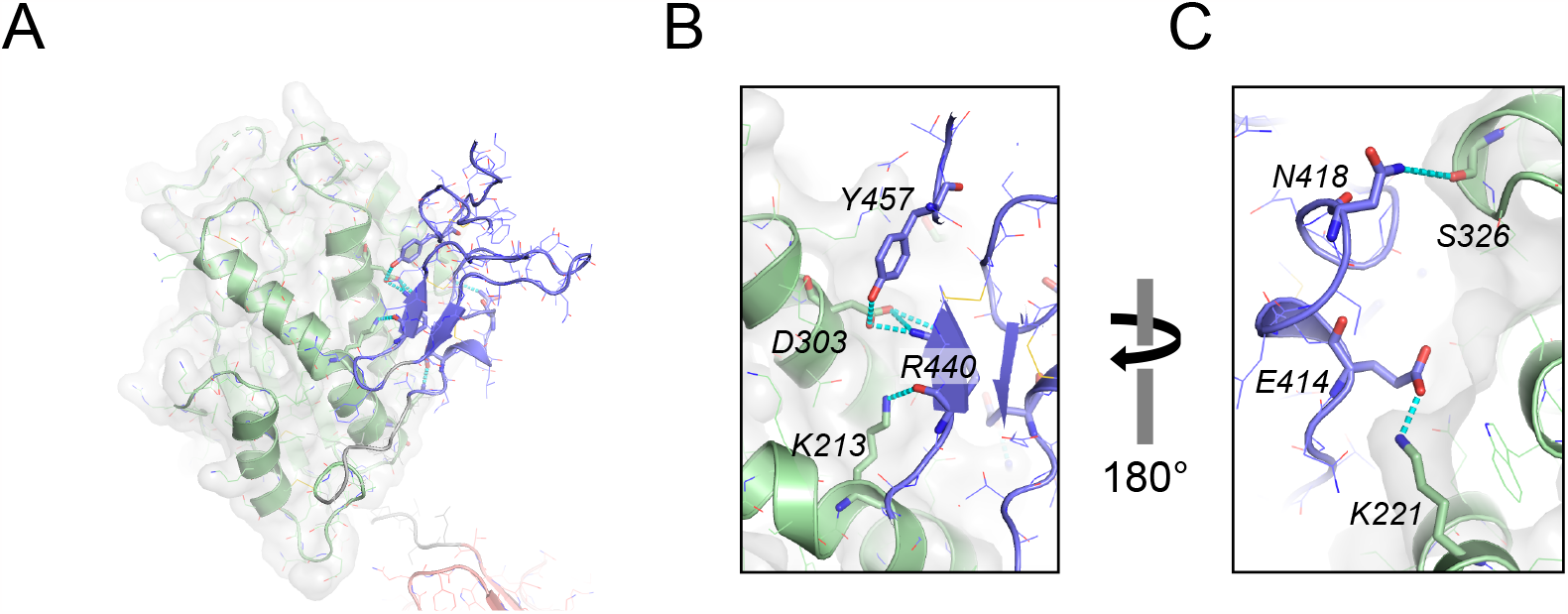
Domain interaction between the TRX and the EGF-like domain. **A**, The overall structure showing the interaction between the two domains. **B** and **C**, Detailed hydrogen (H)-bond interactions are shown. Panel **A** and **B** are presented in the same view. Panel **C** provides a back side view of panel **A** and **B**. The TRX domain and EGF-like domains are colored green and blue, respectively, and the hydrogen bonds are colored cyan. The key residues are depicted as sticks.

***The PA domain*** is composed of approximately 120 residues in length and organized as a central β-barrel with nine strands in two β-sheets, and two α-helices that are located at the periphery (**Suppl. Fig. 1**). Similarly shaped PA domains are known to exist in several protease classes such as subtilases, aminopeptidases, bacterial endopeptidases, as well as two families of sorting receptors, RMRs and VSRs (Mahon and Bateman, 2000; Luo and Hofmann, 2001). However, the exact function of the PA domain has been examined in only a handful of proteins and speculated that the PA domain may serve as a protease-interacting or -regulating domain based on multiple studies (Luo and Hofmann, 2001). Previous structural studies revealed that the PA domain of VSR1 harbors the binding sites for ctVSD and a portion of ssVSD ((Shimada et al., 1997; Tsao et al., 2022; Luo et al., 2014). The shallow pocket is established between β_4_-α_3_ loop and α_4_-α_5_ loop of PA domain where α_4_ unfurls and its residues are pushed away by the bound peptide ((Shimada et al., 1997; Tsao et al., 2022; Luo et al., 2014). The residues between β_5_ and α_5_ (Ser120-Asp137) appear highly flexible as they are disordered in our crystal structure (**Suppl Fig. 1 and 2**). A portion of the corresponding region is also reported to be disordered in the peptide-bound states ((Luo et al., 2014); however, the same portion is fully structured in the apo-form (PDB ID 4TJV) due to stabilization through interaction with the C-terminus and crystallographic contacts. Apart from this C-terminal linker, the PA domain of our AtVSR1 structure overlaps well with the previous apo-form PA domain structure (PDB ID 4TJV), with a root-mean-square-deviation (RMSD) of 1.22 Å over 148 Cα positions of the residues from 26 to 173.

***The TRX domain*** of our AtVSR1 structure reveals four internal disulfide bonds plus an above-mentioned additional disulfide bond between Cys393 and Cys405, which connects it to the EGF-like domain. Although there is no direct interaction between the intramolecular PA and TRX domains, the two domains display extensive interactions with those domains of two neighboring molecules related by a crystallographic 3-fold symmetry, forming a cyclical domain-swapping (**Fig. 3A**). This homotrimer occurs without any conformational change in the PA domain, evidenced by the average RMSD of 0.64 Å between our structure and the structures of the PA domain alone (PDB ID 4TJV, 4TJX, and 8HYG).

**Figure 3.**
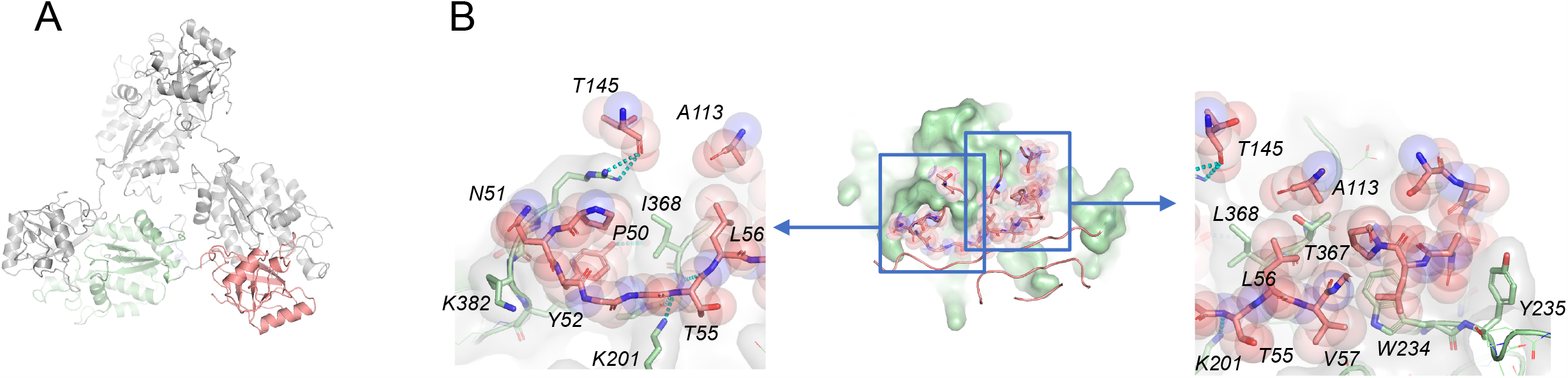
The crystallographic structure exhibits 3-fold symmetry through the swapping of the PA and TRX domains (A). **B**, The detailed interaction between the two domains involves extensive hydrophobic interactions and tight van-der-Waals complementarity. The overall interface measures approximately 1040 Å^2^. The PA domain residues responsible for the interactions are labeled and presented as sticks with transparent spheres. **C**, The hydrophobic surface residues in the TRX domain that participate in the interaction are shown. The PA and TRX domains are colored salmon and green, respectively.

### Alphafold2 model

The full length model of AtVSR1, generated by Alphafold2 (AF2), has been deposited in the AlphaFold protein structure database with an accession number AF-93026-F1 (Jumper et al., 2021). Most of the regions in the model have per-residue-confidence-scores (predicted Local Distance Difference Test, pLDDT) of 0.9 or higher, indicating that the model is highly reliable. Each domain of the model superimposes well with that of the crystal structure (**Suppl Fig. 3A**). However, the interdomain interactions of the AF2 model are completely different from those observed in our crystal structure (**Suppl Fig. 3B and 3C**). The inaccuracy of quaternary structure prediction is a well-known limitation of AF2 (Gao et al., 2022; McCafferty et al., 2023). Despite the caveat, the AF2 model structure still offers valuable insight into how the regions not observed in the crystal structure may look like.

### The oligomerization of VSR1

PDBePISA analysis of the crystallographic *intermolecular* interaction between the PA and the TRX domains reveals that the interface score is 1.0, indicating that VSR1 is very likely to form a trimer in solution (Krissinel and Henrick, 2007). The total buried surface area between the PA and the TRX domains is approximately 1,040 Å^2^, and the interface is formed by 24.6% and 14% of the residues from the PA domain and the TRX domain, respectively. In comparison, the intramolecular interaction between the TRX and EGF-like domains has a PISA interface score of 0, indicating the observed interaction is not significant. The intermolecular interaction between the PA and the TRX domains displays notable hydrophobic interactions mediated by both ends of the loops connected to β3 of the PA domain (**Fig. 3B**). At one end, the residue Tyr52 of PA domain is nestled in a hydrophobic pocket in TRX domain established by Pro370, Thr371, Leu372, Tyr379, Gly381, and Leu383. In addition, nearby hydrophobic residues, Pro50 and Trp175, packed with the phenolic sidechain of Tyr52, forming a larger hydrophobic group. The hydroxyl group of Tyr52 also forms a H-bond with the main chain carbonyl oxygen of Pro370 (**Fig. 3C**). On the other end, a loop formed by Pro84, Gly85, and Arg86 is nested within a valley fabricated by Trp234, Tyr235, Glu353, Gln354, and Ile358. Nearby Thr55 establishes two H-bonds with Leu369 through the main chain amides (**Fig. 3C**). Considering the resolved crystal structures of the TRX domain, the previously hypothesized *intramolecular* PA and TRX domain interaction through the same interface appears unlikely. The linker between the PA and the TRX domains is composed of 7 residues with approximately 26 Å in length. Even when we expanded the linker boundary in our modeling attempt, the proposed intramolecular interaction between the PA and the TRX domains was not possible without disrupting the adjacent residues and dihedral angle violations in linker residues (**Suppl Fig. 4**). This further supports the possibility of an intermolecular interaction that results in a domain-swapped trimer *in vivo*.

### The EGF-like domain

It is noticeable that there is no significant intermolecular interaction by the EGF-like domains in the crystal lattice (**Suppl Fig. 5**). The two additional EGF-like domains that are tandemly located at the C-terminus are not visible in the crystal structure, indicating their high flexibility. It is likely that the string of EGF-like domains is oriented towards the ER membrane and provides flexibility for the PA and TRX domains, enabling them to promote intermolecular interactions (**Suppl. Fig. 3A**). The interaction between the TRX and EGF-like domains is held by multiple H-bond interactions. There are 4 residues each from the TRX domains and the EGF-like domains involved in this interaction. Together, they form 7 H-bonds, out of which 4 are salt bridges (**Fig. 2, Suppl. Table 1**).

## Discussion

### Trimeric nature of AtVSR1

Cao *et al*. conducted elaborate experiments digesting the luminal domain of AtVSR1 to define three protease-resistant domains and their involvement in ssVSD binding. Although the proposed multi-domain nature of AtVSR1 was accurate, interpreting the results based on the estimated molecular weights by SDS-PAGE has some drawbacks (Cao et al., 2000). In the report, they suggested the luminal domain of AtVSR1 could be monomeric in solution based on its elution profile being similar to bovine serum albumin (BSA) on a gel filtration chromatography (Cao et al., 2000). To have similar hydrodynamics to BSA, VSR1 must be tightly packed, suggesting that it could exist as a monomer with an intramolecular interaction between the PA and the TRX domains. In addition, all three EGF-like domains should be tightly packed with the other domains in the same molecule. However, our crystal structure shows that the AtVSR1 has an extended conformation with significant dynamic flexibility of the EGF-like domains. Significantly, the same chromatographic profile of Cao *et al*. also displayed a shoulder peak with a high molecular weight aggregate, which could be attributed to AtVSR1 multimers.

Evidence of AtVSR1 forming a trimer was reported by Kim *et al*. where hemagglutinin-tagged AtVSR1 (AtVSR1:HA) is expressed in transgenic plants, followed by fractionation of the protein extract using Superdex 200 HR 10/30, and identified the existence of 240KDa and ∼80-100KDa species that responded to anti-HA antibody (Ab) (Kim et al., 2010). Subsequent experiments confirmed the 240KDa species to be homopolymer of VSR1. They further dissected the regions responsible for homotrimer formation and concluded that both the transmembrane and C-terminal cytosolic domains were necessary, while the luminal domain was not. This conclusion was based on a series of co-immunoprecipitation assays using deletion and substitution mutants of VSR1. It is possible that the detergents contained in the assay buffer were not suitable for the oligomerization of the luminal domain. Nevertheless, VSR1 with the C-terminal cytosolic domain mutants that cannot form oligomers suffer in vacuolar trafficking efficiency due to localization to the Golgi apparatus instead of the prevacuolar compartment.

Considering the structural evidence of homotrimer formation through the luminal PA and TRX domains presented in full-length luminal domain of AtVSR, the formation of the oligomer through the transmembrane and the flexible C-terminal EGF-like domains might be enhanced by the homotrimer of the luminal domain.

Domain-swapping among homopolymers as observed in our AtVSR1 is not uncommon, in which proteins establish a dimer, a cyclic multimer, or an open-ended aggregate (Kundu and Jernigan, 2004; Bennett, Choe and Eisenberg, 1994a). The swapping can be performed via exchanging a secondary structural element or an intact domain of the participating monomeric subunit. Diphtheria toxin is a classic example that forms a dimer through the latter method (Bennett, Choe and Eisenberg, 1994b). and the conformation in our AtVSR1 structure also swaps domains similarly. The domain swapping that forms a homotrimer in our crystal structure of AtVSR1 is unique. This is because the only viable option for AtVSR1 is the formation of homotrimer (or possibly homomultimer) through *intermolecular* interaction as opposed to previously proposed monomer through *intramolecular* domain interaction, which is exemplified in the diphtheria toxin monomer.

The presence of restrictive linker residues prevents the existence of a functional monomer with the compacted PA-TRX domains. Additionally, due to its cyclic domain swapping, AtVSR1 may also form additional oligomers.

Further analysis of VSR1 in solution using size exclusion chromatography-multi-angle light scattering (SEC-MALS) or sedimentation equilibrium (SE) analysis is necessary to clarify the oligomeric state in solution and the functional implications of oligomerization in the luminal domain.

### VSD-recognition mechanism of AtVSR

The previous structural study by Luo *et al*. showed that a part of the loop between β5 and β6 in the PA domain is displaced by an ssVSD peptide containing an NPIR motif ((Luo et al., 2014). Unexpectedly, this complex structure showed that the interaction between PA domain and a ssVSD peptide is not mediated by the NPIR motif, but rather by three preceding residues of the NPIR motif. Although sensitivity to mutation of the penultimate residue before the NPIR motif was shown by pull-down and other biological assays, a detailed structural analysis showed a sign of promiscuity in the ssVSD peptide binding, as little side chain involvement was observed.

The promiscuity of interaction is evidenced by another structural study by Tsao *et al*., where the authors found that a C-terminal VSD (ctVSD) peptide from CRU1 binds to the VSR1 PA domain in the same way as the ssVSD NPIR-peptide barley aleurain. ctVSD-peptides are composed of four C-terminal residues without a known consensus (Tsao et al., 2022). The structural study revealed a favored tendency; namely, a basic residue is preferred in position 1, and hydrophobic residues (except for proline) are favored in the three remaining positions. Intriguingly, the C-terminal carboxyl group of ctVSD is recognized by Arg95 of PA domain, which is conserved throughout the VSR isoforms. Therefore, the binding site of VSR appears to be attuned to a C-terminal region rather than an internal sequence motif.

Based on the crystal structure of PA domain, Luo *et al*. hypothesized that the *internal* NPIR peptide recognition by VSR1 is established by two separate domains: the three residues preceding the NPIR-binding motif in the PA domain and the NPIR-binding motif in the TRX domain (Luo et al., 2014). As a mechanism to bring two motifs together of two separated domains from a same VSR1 molecule, the authors pointed out the 180° swing motion triggered by the three-residue peptide binding to the PA domain (Luo et al., 2014).

Being estimated from our crystal structure of the luminal domain, the shortest distance between the ctVSD-binding site of the PA domain and the either intra- or inter-molecular TRX domain is approximately 30 Å, which is too far for an NPIR peptide to bind (**Suppl. Fig. 6**). If the domain-swapped trimer presented here exists *in vivo* and the NPIR-binding site is located in the TRX domain, it is still unlikely that the ctVSD-binding site at the PA domain is involved in ssVSD recognition.

### The plausible function of the TRX domain

The TRX domain observed in our AtVSR1 structure shows a typical fold observed among protein disulfide isomerase, PDI-fold (Martin, 1995), featuring a four-helix bundle attached between the classic TRX fold of β_1_α_1_β_2_and β_3_β_4_α_2_ topologies (**Suppl. Fig 1A and 7A**). As noticed in many PDIs, β_4_-strand in the TRX domain of VSR1 is not prominent. The helical bundle observed in the TRX domain of VSR1 has two disulfide bonds; one between helices α1 and α4, and another between helices α2 and α3. There is a 40-residue-long insertion unique to the TRX domain between β_1_α_1_β_2_ and the helical bundle, which features an α-helix and a 3_10_ helix with two disulfide bonds (**Suppl. Fig 7A**). The archetypal TRX-fold containing PDI enzymes have a dithiol CxxC active site at the beginning of α_1_ to catalyze the reduction of a disulfide bond. Superposition of the TRX domain with the PDBs retrieved by the DALI search shows that the dithiol CxxC motif is structurally conserved among those PDIs, even though AtVSR1 has an additional residue between the two conserved cysteines as ^198^CxxxC^202^ (**Suppl. Fig 7B**).

Some of the proteins that are sorted by VSR1 contain disulfide bonds. For example, cruciferin 1 and 3, which are 12S globulins in Arabidopsis, have two and one disulfide bond, respectively. Additionally, aleurain also has two disulfide bonds. Thus it is tempting to speculate that VSR1 engages in the quality control of the cargo proteins with disulfide before or during the sorting process. Furthermore, the formation of vacuoles in seed storage is a rapid process, which is typically completed within 24 to 48 hours (Cao, Duncan and Millar, 2022). Rapid synthesis of the storage proteins may compromise protein folding, and the TRX domain of VSR1 may function as a secondary mechanism to ensure folding of proteins with proper disulfide bond configurations.

An intriguing aspect of TRX domains in general is their ability to recognize protein substrates. Whether the TRX domain of VSR1 is an active enzyme or nature is simply repurposing the fold (Chothia, 1992) for target (ssVSD) binding needs further investigation.

### Ca^2+^ and pH dependence of the EGF-like domain

Ca^2+^-coordination property of the EGF-like domains is achieved by 5 residues, 3 of which through conserved side-chain carboxyl/carboxamide and 2 through the main-chain carbonyls (Handford et al., 1991; Knott et al., 1996; Wouters et al., 2005). Comparison of the EGF-like domain of AVSR1 to known Ca^2+^-binding EGF-like domains show those residue positions are conserved, suggesting the EGF-like domain of vSR1 is likely to bind a Ca^2+^ (**Suppl. Fig. 8**). Deleting the EGF-like domain in PV72, which is a homolog of AtVSR1 from pumpkin seed, displayed a 10-fold decreased affinity for the NPIR peptide compared to the full length. Furthermore, the binding of the NPIR peptide of PV72 is dependent on Ca^2+^, supporting the significance of Ca^2+^-binding motif (Watanabe et al., 2004).

Multiple studies have consistently shown pH-dependent cargo binding and unloading of VSRs. In a recent study, it was demonstrated that the NHX5 and NHX6 antiporters play a critical role in maintaining proper pH homeostasis in vacuoles, and VSR-cargo binding and trafficking rely on pH homeostasis maintained by the two vesicular antiporters (Reguera et al., 2015). The findings further highlight the importance of pH-dependent receptor-cargo interactions in protein trafficking and provide insights into the role of NHX antiporters in regulating VSR function.

Noticeably, the interaction between the TRX and EGF-like domains is held together by 6 H-bonds, 3 of which are salt bridges (**Suppl. Table 1**). The *pK*_*a*_ values of Glu and Asp are 4.4 and 4.0, respectively. Therefore, at low pH, the strength of the salt bridges is likely to be reduced significantly. Likewise, binding of Ca^2+^ to the EGF-like domain is expected to cause a conformational change in the domain, which may influence the interaction with the TRX domain. Further study is needed to fully understand the involvement of the TRX and EGF-like domains, as well as the role of the salt bridges involved, in NPIR peptide binding.

## Conclusion

Decades ago, when we embarked on determining the structure of AtVSR1 generated from the insect cell expression system, we anticipated that it would provide answers to questions regarding the protein’s function. The structure of the full-length luminal domain of AtVSR1 with glycosylation provides fascinating insights into its function. It also raises more intriguing questions that require further investigation. Based on our current structural study, we argue whether the proposed model by Luo *et al*. accurately represents the physiological condition (Luo et al., 2014), that is whether the ctVSD and ssVSD sites are in the PA and TRX domain in tandem at AtVSR1. We instead speculate that the ssVSD binding might be present near the TRX and EGF-like domain interface. This is due to the fact that ssVSD binding is affected by pH and Ca^2+^ ion, and the interaction between the TRX and EGF-like domain is dependent on multiple H-bonds. The Ca^2+^ binding site of the TRX domain may serve as a pH/Ca^2+^ sensor. Further research is needed to elucidate the specific region responsible for ssVSD binding in AtVSR1. Furthermore, whether the TRX domain of AtVSR1 functions as a disulfide bond isomerase needs to be investigated. The PFAM search did not find any other examples of a receptor protein having a chaperone domain. The VSRs may be a truly unique receptor system if the TRX domain functions as disulfide bond isomerase. Given the rapid nature of vacuolar sorting in cells, it is possible that quality control is not given high priority. In this regard, VSRs may serve a dual role of binding cargo and also playing a catch-up function of cargo refolding during the sorting process. Future research should investigate this unique plant system to enhance our understanding of the perspective it offers to all life forms.

## Methods

### Protein expression and purification

Information regarding cloning and protein production can be found in the literature (Rogers et al., 2004; Cao et al., 2000; Paris et al., 1997). In brief, the plasmid containing the luminal domain of VSR1 with a C-terminal 6XHis-tag was transfected into Drosophila S2 cells with the Drosophila Expression System kit. Stably transformed cell lines were selected with hygromycin and transferred to serum-free medium over a month period. Expression of the recombinant proteins was induced with 500 mM copper sulfate for 72 hrs. VSR1 was purified from S2 cell medium using His affinity column followed by a proaleurain peptide-affinity column.

### Crystallography

The crystallization and data collection were disclosed by Rogers *et al. (Rogers et al*., *2004)*. The dataset diffracted to 3.5 Å Bragg spacings was successfully phased by molecular replacement using the PA domain of AtVSR1 (PDB ID 4TJV) as a search model (McCoy et al., 2007). The crystal contained one molecule in the asymmetric unit. The top solution of the MR search had a translation function z-score (TFZ) of 32.6, and the initial phase map showed boundaries for two additional domains as having positive density, all leading to the conclusion that the MR solution was correct. The additional model building was done with Coot, and the refinements were done in Phenix-refine (Emsley et al., 2010; Liebschner et al., 2019). Even at 3.4 Å resolution, most side chain electron densities were legible, and therefore the residues were placed in position. The final structure had a total of 423 residues with N-glycosylation at Asn289. R_work_ and R_free_ were 19.5%, and 23.5%, respectively. Regularization of the coordinate for structural analysis was done in the Maestro protein preparation workflow (Schrödinger, LLC). Structural figures were created with PyMol (Schrödinger, LLC). The coordinate and the reflection file were deposited. The data processing and refinement statistics are presented in (**Suppl. Table 2)**.

## Acknowledgment

We thank the staff at the SSRL beamline for synchrotron data collection. We would like to express sincere gratitude to Dr. John C. Rogers and Sally W. Rogers for their initial work for the study and providing purified AtVSR1, and Dr. Marisa Otegui for her insightful discussion of the manuscript. Original research was supported by NSF (MCB-2043248), USDA-NIFA (2023-67013-39629, 2003-35318-13672).

## Author Contributions

**HP** collected and analyzed data, and wrote manuscript. **BY** prepared crystals and analyzed data. **DJP** analyzed data and wrote manuscript. **SVP** analyzed data. **CK** initiated the project, secured funding, analyzed data, and wrote manuscript.

